# Loss of age-accumulated *crh-1* circRNAs ameliorate amyloid β-induced toxicity in a *C. elegans* model for Alzheimer’s disease

**DOI:** 10.1101/2024.04.09.588761

**Authors:** Hussam Alshareef, Thomas Ballinger, Everett Rojas, Alexander M. van der Linden

## Abstract

Circular RNAs (circRNAs) are non-coding RNAs mostly derived from exons of protein-coding genes via a back-splicing process. The expression of hundreds of circRNAs accumulates during healthy aging and is associated with Alzheimer’s disease (AD), characterized by the accumulation of amyloid-beta (Aβ) proteins. In *C. elegans*, many circRNAs were previously found to accumulate during aging, with loss of age-accumulated circRNAs derived from the CREB gene (circ-*crh-1*) to increase mean lifespan. Here, we used *C. elegans* to study the effects of age-accumulated circRNAs on the age-related onset of Aβ-toxicity. We found that circ-*crh-1* mutations delayed Aβ-induced muscle paralysis and lifespan phenotypes in a transgenic *C. elegans* strain expressing a full-length human Aβ-peptide (Aβ_1-42_) selectively in muscle cells (GMC101). The delayed Aβ phenotypic defects were associated with inhibiting the deposition of Aβ aggregates, and thus, genetic removal of circ-*crh-1* provides protection against Aβ-induced toxicity. Consistent with a detrimental role for age-accumulated circRNAs in AD, circ-*crh-1* expression level is elevated after induction of Aβ during aging, whereas linear *crh-1* mRNA expression remains unchanged. Finally, we show that a circ-*crh-1* upregulated collagen gene, *col-49*, promotes Aβ-induced paralysis. Taken together, our results show that the loss of an age-accumulated circRNA exerts a protective role on Aβ-induced toxicity, demonstrating the utility of *C. elegans* for studying circRNAs in AD and its relationship to aging.

## INTRODUCTION

Circular RNAs (circRNAs) have emerged as an intriguing class of non-coding RNAs with unique closed-loop structures. CircRNAs are generated through a process known as back-splicing during conventional RNA splicing, where the 3’ and 5’ ends of a pre-mRNA molecule are covalently bonded, yielding a circular configuration (Li et al., 2018). Most identified circRNAs are produced from exons of protein-coding genes (Zhang et al., 2014). Their lack of free ends confer circRNAs a degree of resistance to exoribonuclease digestion compared to their linear counterparts (Jeck et al., 2013), which can contribute to their stability and abundance. Despite these characteristics, the functions of many circRNAs remain largely elusive, although their functions appear to be intertwined with the molecules they interact with.

CircRNAs accumulate during the normal process of aging in *C. elegans* (Cortés-López et al., 2018)*, Drosophila* (Hall et al., 2017; Westholm et al., 2014), and mice (Gruner et al., 2016) and are found to play both positive and negative roles in the aging of *C. elegans*, *Drosophila*, and various mammalian tissues (Kim et al., 2021; Knupp and Miura, 2018). Prominent examples include the extension of lifespan through *circSfl* transgenic overexpression in *Drosophila* (Weigelt et al., 2020) and the loss of the circRNA derived from the host gene *crh-1*/CREB in *C. elegans* (Knupp et al., 2022).

Additionally, growing evidence suggests key roles of circRNAs in Alzheimer’s disease (AD) by affecting mechanisms such as neuroinflammation, oxidative stress and autophagy, as well as amyloid-beta (Aβ) production and degradation (Beylerli et al., 2024). For example, circular RNA *ciRS-7* (also known as *CDR1as*) inhibits the activity of microRNA, *mir-7* (Hansen et al., 2013), which subsequently effects the accumulation of Aβ plaques in AD (Shi et al., 2017; Sun et al., 2023). Considering that brain aging is highly associated with AD pathogenesis, circRNAs that accumulate with aging may also contribute to AD. These circRNAs could act as molecular sponges, sequestering microRNAs (miRNAs) (Hansen et al., 2013; Zhang et al., 2020) and RNA-binding proteins (RBPs) (Chen, 2020; Patop et al., 2019) away from their messenger RNA (mRNA) targets, thereby altering the splicing or expression patterns of these mRNAs.

However, our understanding of the functional roles of age-accumulated circRNAs in AD remains limited. *C. elegans* presents a powerful model organism for studying age-associated circRNAs in AD. Previously, we demonstrated that a majority of circRNAs expressed in *C. elegans* accumulate during aging (Cortés-López et al., 2018). Using CRISPR/Cas9, we genetically removed two abundant age-accumulated circRNAs derived from the *crh-1* gene (circ-*crh-1*) encoding the homolog of CREB without disrupting the linear RNA and its associated activated protein (Knupp et al., 2022). Genetic loss of this age-accumulated circ-*crh-1* extended the mean lifespan of *C. elegans* (Knupp et al., 2022), suggesting that circ-*crh-1* abundance might contribute to age-related decline. Here, we extended our findings by testing the impact of circ-*crh-1* removal on a severe model of inducible amyloidosis in *C. elegans*. We used the transgenic *C. elegans* strain (GMC101) expressing human Aβ_1-42_ peptides constitutively in muscle cells that mimics the pathological features of AD (McColl et al., 2012). We showed that loss of circ-*crh-1* expression delayed the Aβ-induced paralysis of GMC101, improved its reduced lifespan, and reduced the number of Aβ aggregates. Moreover, we find that the predicted collagen gene, *col-49*, is upregulated in circ-*crh-1*(-) mutants and contributes to Aβ-induced paralysis. Together, our results show that the loss of an abundant and age-accumulated circRNA ameliorates Aβ-induced toxicity in a *C. elegans* model for AD.

## RESULTS

We previously demonstrated that expression of circ-*crh-1* accumulates with *C. elegans* aging (Cortés-López et al., 2018) and that loss of circ-*crh-1* expression results in a significant extension of mean lifespan (Knupp et al., 2022). To directly test whether circ-*crh-1* expression plays a role in Aβ-induced toxicity, we used the GMC101 strain (further referred as *unc-54p::A*β*_1-42_*) in which expression of human Aβ_1-42_ peptides constitutively in muscle cells provokes an inducible age-progressive full-body paralysis when shifted from the non-restrictive 20°C temperature to a higher 25°C temperature (McColl et al., 2012). We found that the full body paralysis phenotype of *unc-54p::A*β*_1-42_* animals is ameliorated in two independent mutant alleles of circ-*crh-1* (*syb385* and *syb2657*) that exhibit a complete loss of circ-*crh-1* expression but show normal linear *crh-1* expression (**Figure 1A**; (Knupp et al., 2022). Thus, loss of circ-*crh-1* expression significantly delayed the onset of Aβ-induced paralysis.

**Fig 1:**
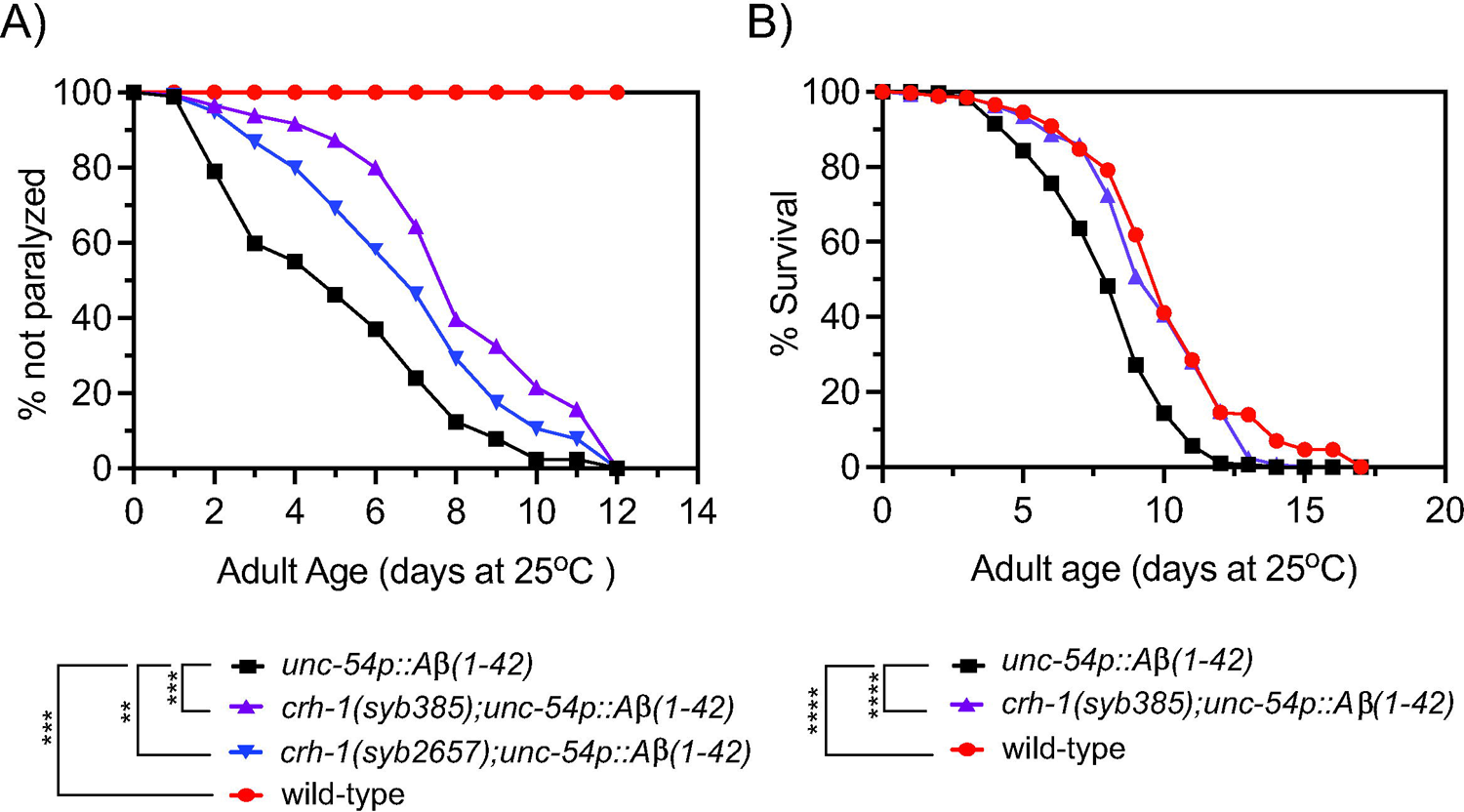
Loss of circ-*crh-1* delays paralysis and decrease the short lifespan of an Aβ-induced proteotoxicity model. (A) The onset of paralysis measured in the Aβ-proteotoxicity model strain, GMC101 (*unc-54p::A*β*_1-42_*) and the GMC101 strain carrying either a *syb385* mutation (purple) or *syb2657* mutation (blue) after a temperature upshift to 25°C. Wild-type (red) or *unc-54p::A*β*_1-42_*(GMC101, black) animals at 20°C do not show paralysis. 3 independent trials with n>140 animals for each assay and genotype in the presence of 0.5 µM FUdR. Asterisks indicate statistical significance with \*\**p*<0.01, \*\*\**p*<0.001. (B) Lifespan curves for *unc-54p::A*β*_1-42_* (GMC101) animals compared to *crh-1(syb385); unc-54p::A*β*_1-42_* and wild-type animals at 25°C. Induction of *unc-54p::A*β*_1-42_* shortens lifespan compared to wild-type (\*\*\*\**p*<0.0001, Mantel-Cox log-rank test), which can be reversed by *syb385* mutations. There is a non-significant difference in mean lifespan between *crh-1(syb385); unc-54p::A*β*_1-42_*and wild-type (*p*<0.035, Mantel-Cox log-rank test). See **Table S3** for lifespan statistics. n=3-4 independent lifespan assays were performed with n=90-150 animals for each assay and genotype in the presence of 0.5 µM FUdR (see Material and Methods).

To further test the protective effect of circ-*crh-1* expression *A*β-induced toxicity, we next tested its loss on the lifespan of *unc-54p::A*β*_1-42_* expressing animals. Previous work has shown that the expression of *A*β*_1-42_* in body wall muscle cells severely decreased lifespan (Gallrein et al., 2021). Similarly, we found that *unc-54p::A*β*_1-42_* (GMC101) expressing animals led to a significantly shorter mean lifespan compared to wild-type controls (20.5% reduction, 8.10 days for GMC101 versus 10.19 days for wild-type, *p*<0.0001) when animals were shifted from the non-restrictive 20°C temperature to the higher 25°C temperature (**Figure 1B**). Interestingly, *crh-1(syb385)* was able to restore the reduced lifespan of *unc-54::A*β*_1-42_*animals back to wild-type levels (9.71 days for *crh-1(syb385)*; *unc-54::A*β*_1-42_* versus 8.10 days for GMC101, *p*<0.0001) (**Figure 1B**). No significant differences were observed in mean lifespan between *unc-54::A*β*_1-42_*and wild-type animals (*p*=0.614) at the non-restrictive 20°C temperature (no Aβ-induction) (**Figure S1**). Thus, circ-*crh-1* mutations prevented lifespan shortening induced by Aβ at 25°C. Together, these findings suggest that loss of circ-*crh-1* expression protects *C. elegans* from Aβ-induced toxicity.

We next examined the impact of Aβ-induction on the expression of the age-accumulated circ*-crh-1* in aged *unc-54::A*β*_1-42_* animals compared to wild-type. We therefore conducted RT-qPCR analysis to measure the RNA fold change of circ-*crh-1* expression (i.e. *cel-circ_0000439*) in 4-day old adults compared to 1-day old adults for wild-type and between *unc-54::A*β*_1-42_* animals at 25°C (after Aβ-induction). As expected, the fold-change ratio of circ-*crh-1* gene expression was higher in *unc-54::A*β*_1-42_*animals than wild-type by 1.25-fold (**Figure 2A**). Importantly, expression of linear *crh-1* was not significantly affected after Aβ-induction in 1-day and 4-day old adults (**Figure 2A**).

**Fig 2:**
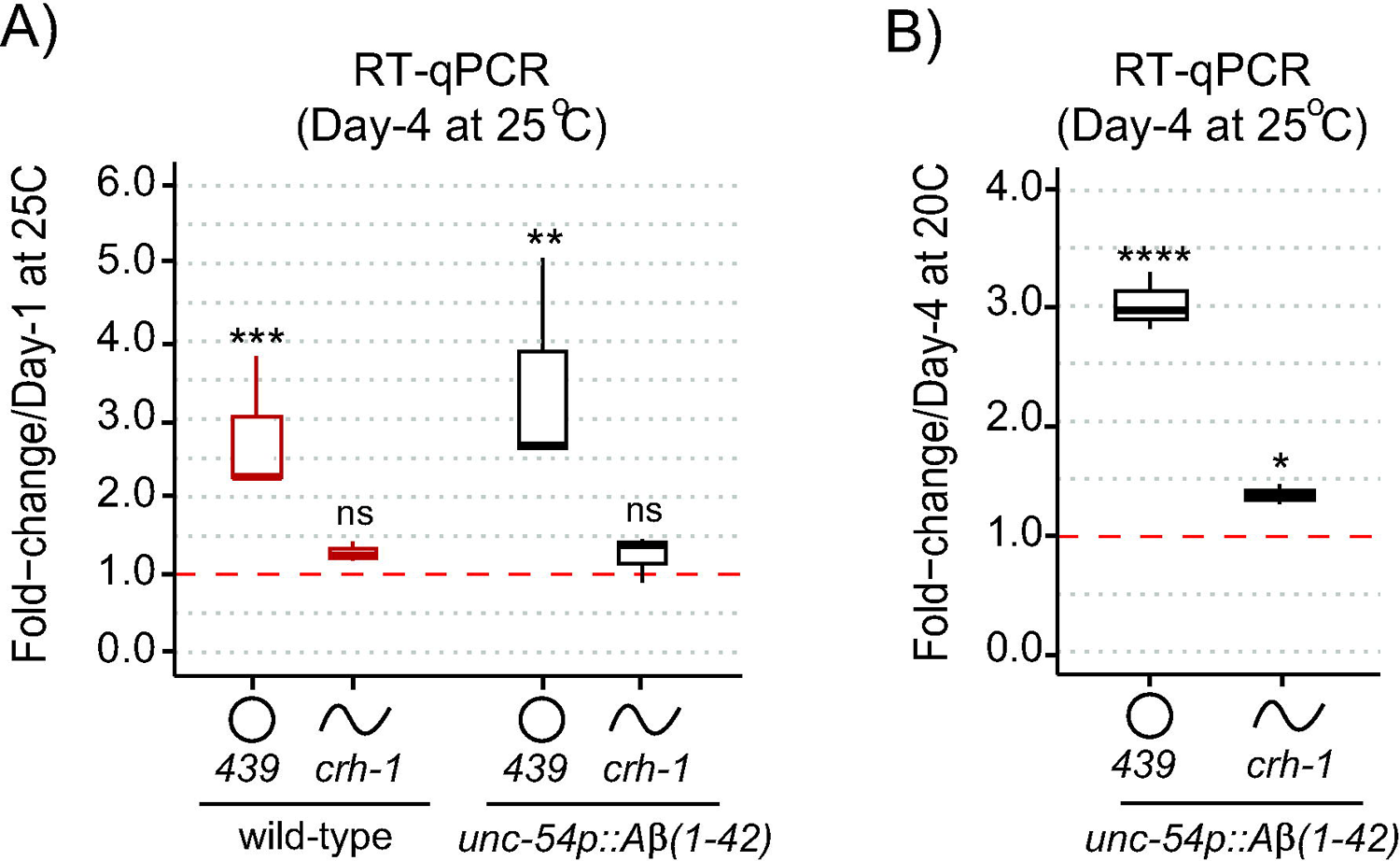
circ-*crh-1* expression is increased after Aβ-induction during aging. (A) RT-qPCR expression of the abundant circular circ-*crh-1* (*cel_circ_0000439*) and linear *crh-1* transcripts in 4-day adults normalized to 1-day adults of wild-type and *unc-54p::A*β*_1-42_* (GMC101) at 25°C. (B) RT-qPCR expression of circular and linear *crh-1* transcripts in day-4 adults of *unc-54p::A*β*_1-42_*(GMC101) at 25°C (after Aβ-induction) normalized to 1-day adults of *unc-54p::A*β*_1-42_* (GMC101) at 20°C (no Aβ-induction). circ-*crh-1* expression increases after induction of Aβ. n=3 independent biological samples for both (A) and (B). For RT-qPCR expression analysis, data in panels (A) and (B) was normalized to *cdc-42* mRNA. Data is represented as mean ± SEM. ns, not significant. \**p*<0.05, \*\**p*<0.01, \*\*\**p*<0.001, \*\*\*\**p*<0.0001.

Similar results were observed when normalizing circ-*crh-1* gene expression of 4-day old *unc-54::A*β*_1-42_* adults at 25°C (after Aβ-induction) to 20°C (before Aβ-induction) (**Figure 2B**). These results suggest that Aβ-induction positively regulates circ*-crh-1* expression, consistent with a detrimental role for age-accumulated circRNAs in AD.

Next, we asked whether loss of circ-*crh-1* expression reduced Aβ-aggregate deposits in *unc-54::A*β*_1-42_* animals, which results in age-progressive paralysis and a shortened lifespan (**Figure 1**). In order to rule out the effects of circ-*crh-1* loss on the transcription of *A*β*_1-42_*expression rather than simply reducing *A*β aggregates in muscle, we first used RT-qPCR analysis to measure Aβ mRNA levels at different aging time-points. We found no significant differences in Aβ gene expression between *unc-54::A*β*_1-42_* and *crh-1(syb385)*; *unc-54::A*β*_1-42_* animals at 1, 2, and 3-day old adulthood when raised at 25°C (**Figure 3A**). We then utilized the sensitive amyloid-binding dye X-34 (Link et al., 2001b) to specifically stain and visualize *in vivo* Aβ aggregates in muscle cells of *C. elegans* as reported (McColl et al., 2012) and assessed whether circ-*crh-1*(-) mutations reduce the number of Aβ aggregates puncta in *unc-54::A*β*_1-42_*expressing worms. Consistent with the delayed onset of the age-progressive Aβ-induced paralysis and restoration of the shortened lifespan, we found that the number of X-34 positive Aβ-aggregates were significantly reduced (*p*<0.05) in *crh-1(syb385)* mutants carrying *unc-54::A*β*_1-42_*compared to GMC101 in 5-day old adults when raised at 25°C (**Figure 3B-C**). Thus, loss of circ-*crh-1* expression inhibits the accumulation of Aβ aggregates.

**Fig 3:**
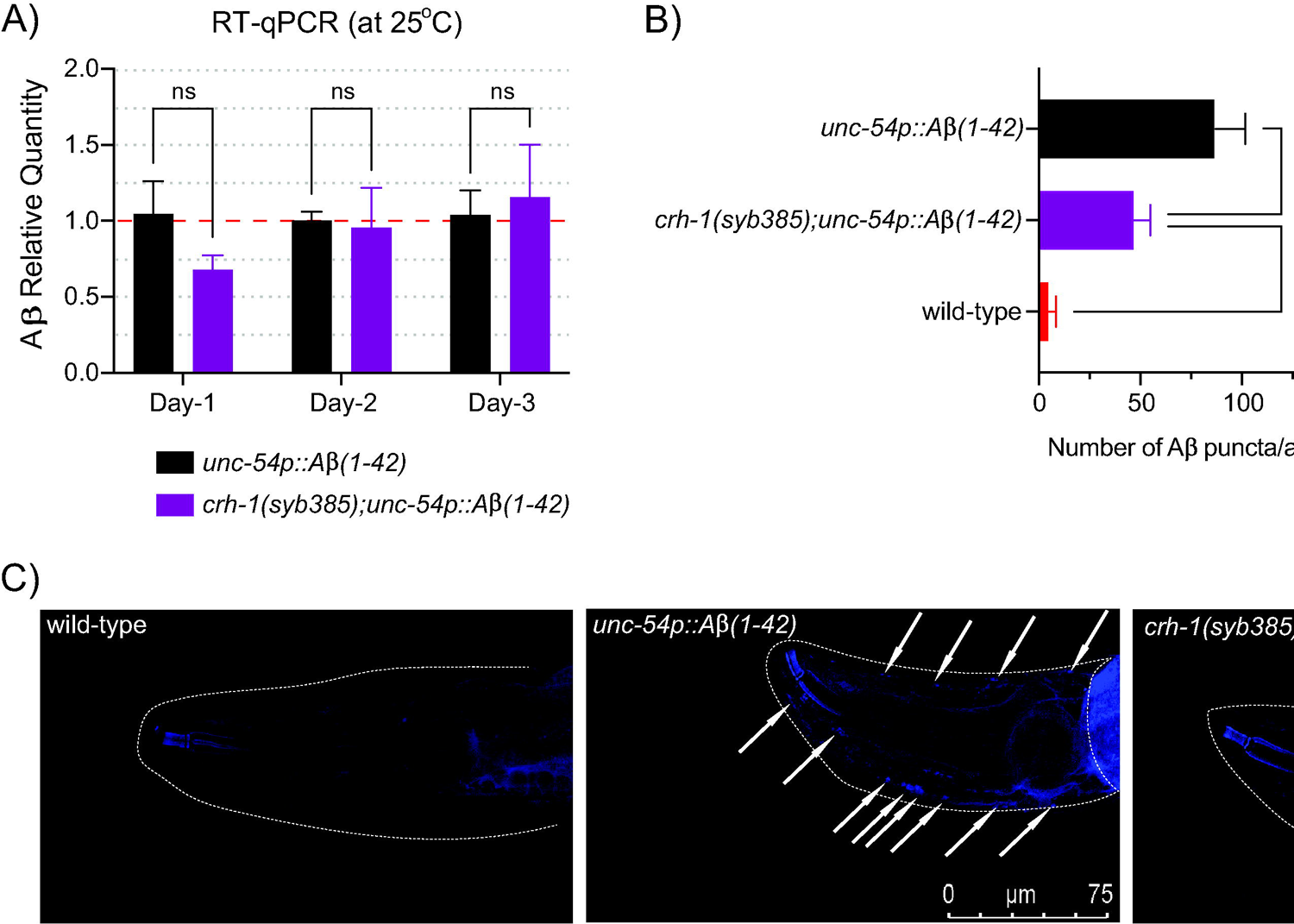
circ-*crh-1* mutation reduces Aβ-aggregation. (A) Relative quantity of Aβ_1-42_ gene expression in *unc-54p::A*β*_1-42_* (GMC101, black), *crh-1(syb385); unc-54p::A*β*_1-42_*(purple) animals during aging (1-day, 2-day, and 3-day old adults at 25°C) as determined by RT-qPCR. Data was normalized to *cdc-42* mRNA. Data is represented as mean ± SEM. n=3 independent biological samples, ns, not significant. (B) Quantitative analysis of Aβ_1-42_ deposits in the head region of wild-type (red), *unc-54p::A*β*_1-42_* (GMC101, black), and *crh-1(syb385); unc-54p::A*β*_1-42_* (purple) animals. The quantity is expressed as mean number ± SEM of Aβ deposits/area of the head region. n=10 animals per genotype. \**p*<0.05, \*\*\*\**p*<0.0001. (C) Representative images of X-34 staining in wild-type (left), *unc-54p::A*β*_1-42_* (GMC101, middle) and *crh-1(syb385); unc-54p::A*β*_1-42_* (right) animals. White arrows indicate Aβ_1-42_ reactive deposits (arrows) in the worm head (dotted white line). Scale barLrepresentsL75 μm.

We next asked whether circ-*crh-1* expression in muscle could restore the delayed onset of Aβ-induced paralysis of *crh-1(syb385)* mutants expressing *A*β*_1-42_* in muscle cells. To investigate this question, we used tissue-specific rescue experiments. We previously showed that circ-*crh-1* expression in neurons is an important determinant for lifespan regulation (Knupp et al., 2022). We decided to create *crh-1(syb385); unc-54::A*β*_1-42_* transgenic animals that express circ-*crh-1* under select tissue-specific promotors. We cloned the *crh-1*(exon 4) circularizing sequence in between the left and right reverse complementary match (RCM) sequences, and used muscle (*myo-3*), pan-neural (*rab-3*), or germline (*pie-1*) specific promoters to drive the circ-*crh-1* expression transgene (**Figure 4A**). Expression of circ-*crh-1* under control of the muscle-specific *myo-3* promoter could partially restore the delayed onset of Aβ-induced paralysis of *crh-1(syb385); unc-54::A*β*_1-42_* (**Figure 4B**), suggesting that circ-*crh-1* expression in muscle is an important determinant for Aβ-induced toxicity. Surprisingly, however, circ-*crh-1* expression driven by *rab-3* and *pie-1* promoters also partially restored Aβ-induced paralysis (**Figure 4B**). These results suggest that in addition to muscle, circ-*crh-1* expression may have additional requirements in other tissues to alter Aβ-induced paralysis.

**Fig. 4:**
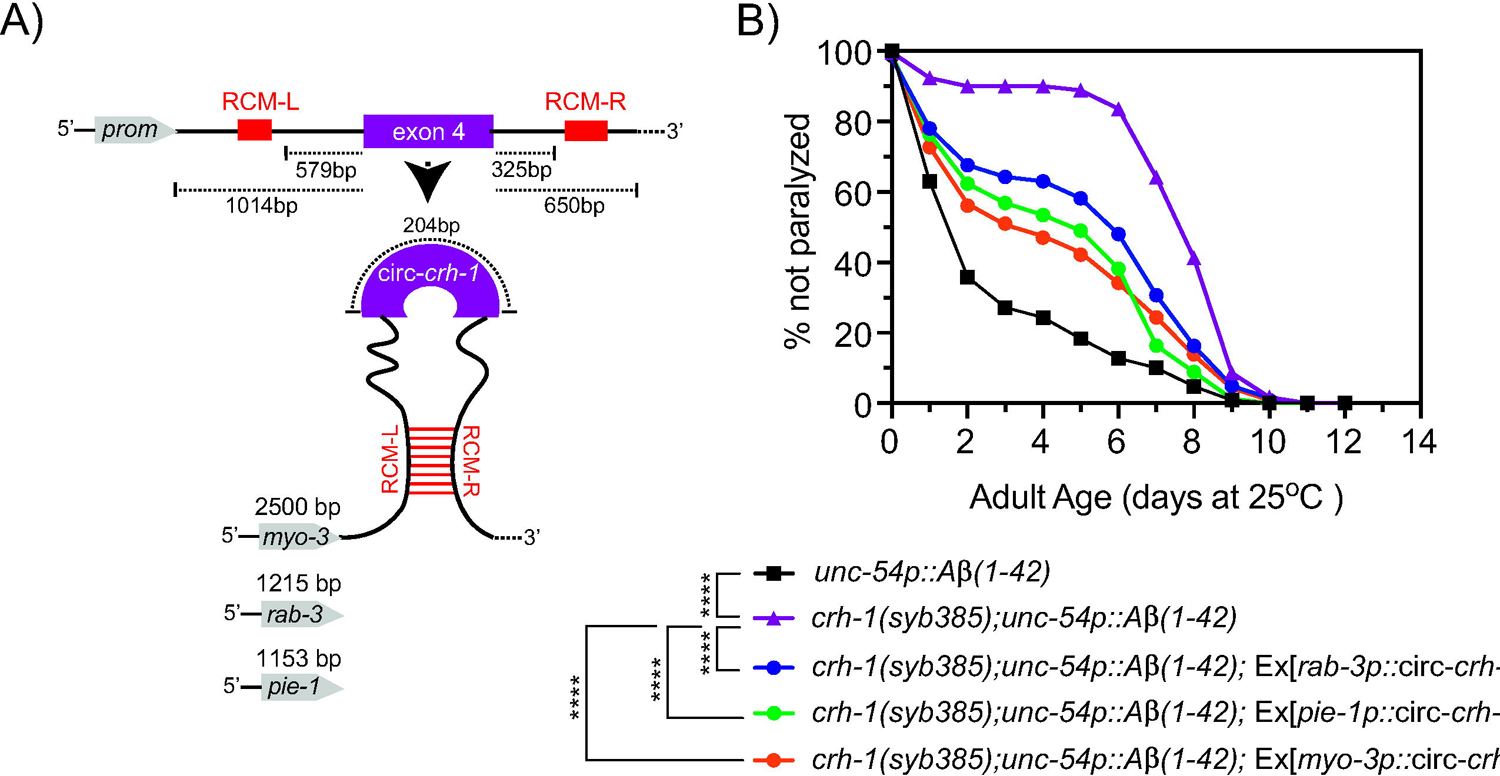
Expression of circ-*crh-1* in multiple tissues can partially rescue the delayed onset of Aβ-induced paralysis in circ-*crh-1* mutants (A) Schematic of plasmid-based minigene used to overexpress circ-*crh-1* under control of tissue-defined promoters. Shown is the *crh-1*(exon4) and reverse complementary match sequences (RCM-L and RCM-R) predicted to facilitate back-splicing of circ-*crh-1*. Promoters and lengths used are *myo-3* with 2,500 bp for muscle expression, *rab-3* with 1,215 bp for pan-neural expression, and *pie-1* with 1,153 bp for germline expression. (B) Paralysis measured in *crh-1(syb385); unc-54p::A*β*_1-42_* animals overexpressing circ-*crh-1* in *myo-3*-expressing muscle cells, *rab-3*-expressing neurons, and *pie-1*-expressing germline cells compared to *unc-54p::A*β*_1-42_* (GMC101, black), *crh-1(syb385); unc-54p::A*β*_1-42_* (purple) animals. 3 independent trials with n>140 animals for each assay in the presence of 0.5 µM FUdR. Asterisks indicate statistical significance with \*\*\**p*<0.001 and \*\*\*\**p*<0.0001.

We previously showed that circ-*crh-1*(-) mutant exhibit widespread transcriptomic changes that might impact various age-related pathways (Cortés-López et al., 2018). Notably, among the identified genes, subset included collagen-encoding genes with many showing upregulation in circ-*crh-1*(-) mutants. Interestingly, cuticular collagens are implicated in Aβ aggregate formation and clearance pathways of Aβ in *C. elegans* (Jongsma et al., 2023). We hypothesized that the upregulation of cuticular collagen gene expression contributes to the amelioration of Aβ-induced toxicity in circ-*crh-1*(-) mutants carrying the *unc-54::A*β*_1-42_* transgene. To test this hypothesis, from the 21 upregulated collagen genes previously identified in *crh-1(syb385)* mutants (Knupp et al., 2022), we selected six cuticular collagen genes of interest, including collagens that have a known association with lifespan such as *col-49* and *col-179* (Palani et al., 2023).

Among the collagen genes tested with RT-qPCR analysis, only *col-49* exhibited a strong upregulation in *crh-1(syb385)* mutants expressing *unc-54::A*β*_1-42_* compared to GMC101 controls at 25°C in 3-day old adults (**Figure 5A**). We next generated *col-49* deletion mutants using a CRISPR/Cas9 strategy and crossed the mutant with GMC101 to assess Aβ-induced toxicity through a paralysis assay at 25°C. We found that *col-49(syb8747); unc-54::A*β*_1-42_*animals exhibited an exacerbated age-progressive paralysis compared to the control (GMC101) (**Figure 5B**). *col-49* mutants without the *unc-54::A*β*_1-42_* transgene did not display paralysis at 25°C (**Figure 5B**). Moreover, no detectable differences were observed in total collagen levels in *crh-1(syb385)* mutants compared to wild-type under different cultivating temperatures (**Figure S2**). Collectively, these findings suggest that the expression of the predicted cuticular collagen gene, *col-49*, is upregulated in circ-*crh-1*(-) mutants and contributes to Aβ-induced paralysis.

**Fig 5:**
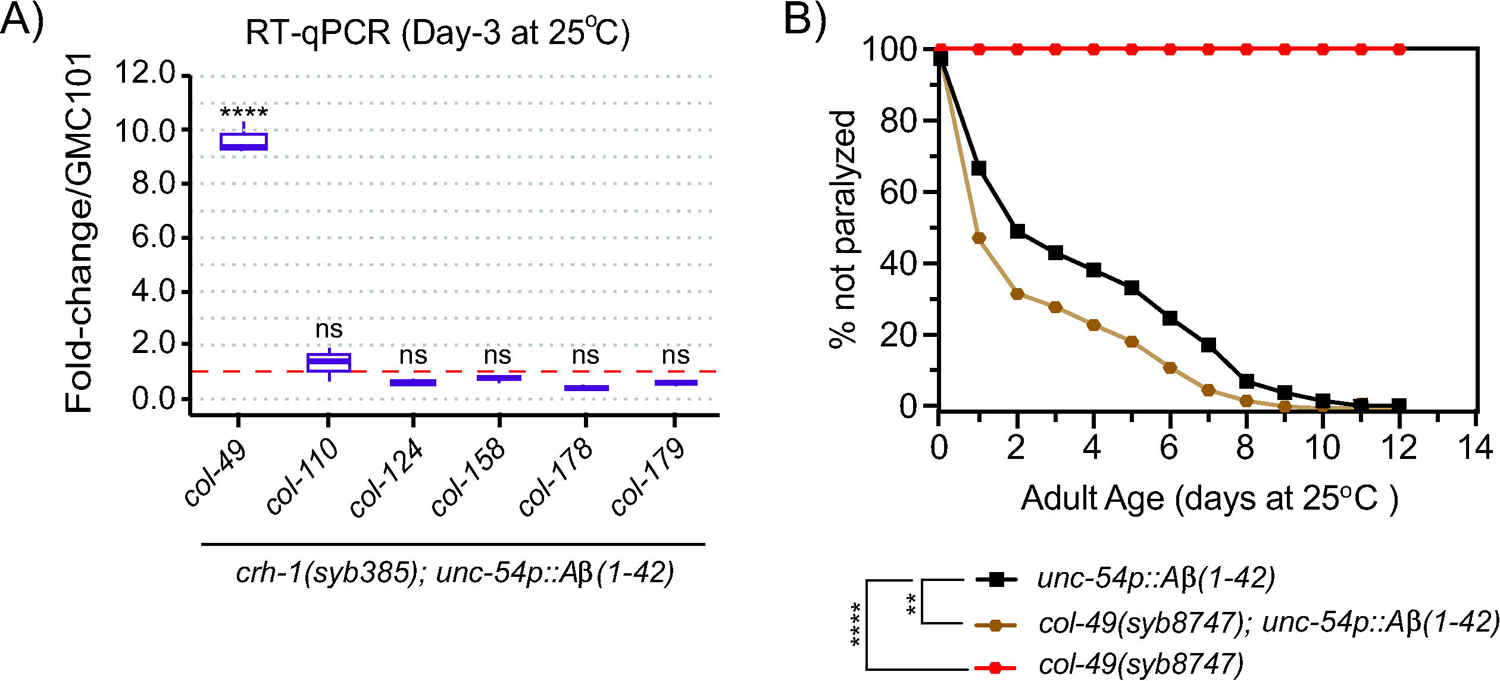
circ-*crh-1* mutations increase *col-49* expression after Aβ-induction, whereas loss of *col-49* promotes Aβ-induced paralysis. (A) RT-qPCR expression of 6 predicted cuticular collagen genes in *crh-1(syb385); unc-54p::A*β*_1-42_* normalized to *unc-54p::A*β*_1-42_* (GMC101) at 25°C (after Aβ-induction) in 3-day old adults. *col-49* expression is strongly increased after Aβ-induction in *crh-1(syb385)* mutants. Data is represented as mean ± SEM and was normalized to *act-1* mRNA. n=3 independent biological samples. ns, not significant. \*\*\*\**p*<0.0001. (B) Paralysis measured in *unc-54p::A*β*_1-42_*animals carrying a *col-49(syb8747)* mutation (brown) compared to *unc-54p::A*β*_1-42_* (GMC101, black) and *col-49(syb8747)* mutant animals (red). The *syb8747* allele has a 1,180 bp deletion generated by CRISPR/Cas9 (see Materials and Methods). 3 independent trials with n>140 animals for each assay in the presence of 0.5 µM FUdR. Asterisks indicate statistical significance with \*\**p*<0.01 and \*\*\*\**p*<0.0001.

## DISCUSSION

The expression of circRNAs accumulates during aging in *C. elegans*, *Drosophila* and mice (Cortés-López et al., 2018; Gruner et al., 2016; Hall et al., 2017; Westholm et al., 2014) as well as in age-associated disorders such as Alzheimer’s disease (AD) (Dube et al., 2019), but the role of age-accumulated circRNAs in AD remains unclear. We assessed whether loss of a single abundant and age-accumulated circRNA, called circ-*crh-1*, could protect against amyloid β-induced toxicity using a well-established *C. elegans* transgenic strain that expresses human Aβ_1-42_ in muscle cells, which results in age-progressive full body paralysis and shortened lifespan. We found a significant delay in the onset of Aβ-induced paralysis in two independent circ-*crh-1*(-) mutants, which could partially be rescued by re-introducing circ-*crh-1* expression in muscle, neurons and germline cells. In addition, we observed that circ-*crh-1*(-) mutants improve the reduced lifespan of muscle expressing Aβ_1-42_ animals. We further demonstrated that genetic removal of circ-*crh-1* expression results in a reduction of Aβ aggregates, suggesting that loss of a single age-accumulated circRNA protects against the age-related onset of Aβ toxicity in *C. elegans*.

Our transgenic experiments implicate circ-*crh-1* expression in muscle by which circ-*crh-1* could delay the onset of Aβ_1-42_-induced paralysis in the GMC101 strain. We also observed that circ-*crh-1* expression in neurons and germline cells could partially restore the delayed onset of Aβ_1-42_-induced paralysis caused by circ-*crh-1* mutations. The GMC101 strain is an inducible model for muscle expressing Aβ_1-42_ aggregation and proteotoxicity (McColl et al., 2012). Aβ peptides have the capability to spread and transfer between cells (Domert et al., 2014). In *C. elegans*, intracellular Aβ_1-42_ peptides expressed in a subset of neurons are able to spread to other cells and distal tissues, and targeted depletion of neuronal Aβ can systemically delay Aβ aggregation (Gallrein et al., 2021). Our rescue experiments suggest that circ-*crh-1* expression is required in other tissues besides muscle for Aβ_1-42_-induced paralysis, but it remains uncertain whether the GMC101 strain exhibits systemic defects and aggregation of Aβ_1-42_ beyond the tissue of expression (i.e. muscle). It might also be possible that overexpression of circ-*crh-1* in neurons and germline cells leads to non-cell-autonomous rescue of muscle expressing Aβ_1-42_ aggregates (Nussbaum-Krammer and Morimoto, 2014). Further experiments demonstrating *in vivo* expression of circ-*crh-1* coupled with labeling Aβ_1-42_ could offer insight into the mechanisms through which circ-*crh-1* regulates Aβ_1-42_ aggregation and its phenotypic consequences.

Using transcriptome-wide analysis, we previously showed an upregulation of multiple collagen-encoding genes in circ-*crh-1*(-) mutants (Knupp et al., 2022). Collagens have previously been linked to Alzheimer’s disease, with several collagens influencing Aβ-aggregate formation (Cheng et al., 2009; Tong et al., 2010). Moreover, a recent study reported that *C. elegans* cuticular collagens are implicated in extracellular Aβ-aggregate formation and clearance (Jongsma et al., 2023). We selected 6 upregulated collagen genes in circ-*crh-1*(-) mutants (Knupp et al., 2022) and tested them in circ-*crh-1*(-) mutants expressing Aβ_1-42_ in muscle. We found that expression levels of the predicted cuticular collagen, *col-49*, with a known role in lifespan regulation (Palani et al., 2023) and cuticular integrity (Jackson et al., 2014) was significantly increased, while the other tested collagen genes were not different from the GMC101 control. We do not yet know how loss of circ-*crh-1* expression results in increased *col-49* mRNA levels in the presence of muscle expressing Aβ_1-42_. We favor the possibility that circ-*crh-1* interacts with RNA-binding proteins (RBPs) to regulate their expression and function by acting as a sponge, decoy, scaffold or recruiter, which could affect the fate of mRNA targets of RBPs through post-transcriptional processes (Chen, 2020; Patop et al., 2019). In this scenario, and consistent with loss of *circ-crh-1* expression resulting in transcriptomic changes (Knupp et al., 2022), circ-*crh-1* may sequester away RBPs from *col-49* mRNA targets, which in turn alters its expression.

Collagen biosynthesis and stability in *C. elegans* can affect Aβ-aggregate levels (Jongsma et al., 2023). While we did not observe any changes in overall collagen levels in whole circ-*crh-1*(-) mutant animals, we found that mutants lacking *col-49* exacerbate the Aβ_1-42_-induced paralysis. This exacerbation might be explained by circ-*crh-1* indirectly modulating *col-49* mRNA levels through one or more RBPs, thereby altering Aβ-induced toxicity. Nevertheless, as the specific RBPs have yet to be identified, it remains formally possible that the exacerbated Aβ_1-42_-induced paralysis observed in *col-49* mutants is not dependent on circ-*crh-1*.

In conclusion, our study shows that circ-*crh-1* expression is important for modulating Aβ-induced toxicity in Alzheimer’s Disease (AD) and could pave the way for using *C. elegans* to study circRNAs in AD and its relationship to aging.

## ACKNOWLEDGEMENTS

We thank members of the van der Linden for useful discussions and feedback on the manuscript. We thank SunyBiotech for help with cloning the *myo-3p::circ-crh-1* construct and creating transgenic lines. We thank the Cellular and Molecular Imaging (CMI) Core facility at the University of Nevada, supported by a grant from the NIH National Institutes of General Medical Sciences (P30 GM145646), for providing equipment and resources. We thank the Caenorhabditis Genetics Center (GCG), funded by the NIH Office of Research Infrastructure Programs (P40 OD010440), for providing strains used in this study.

## AUTHOR CONTRIBUTIONS

HA and AVDL designed the study. HA, TB and ER performed the experiments. HA, TB, ER and AVDL analyzed the data. HA and AVDL wrote the manuscript. AVDL supervised the study.

---

**Fig S1:** Lifespan analysis of GMC101 at 20°C Lifespan curve for *unc-54p::A*β*_1-42_* (GMC101) animals compared to wild-type animals at 20°C. There is a non-significant difference in mean lifespan between *unc-54p::A*β*_1-42_* and wild-type controls (*p*=0.642, Mantel-Cox log-rank test). See **Table S3** for lifespan statistics. n=4 independent lifespan assays were performed with n=90-120 animals for each assay and genotype in the presence of 0.5 µM FUdR (see Material and Methods).

**Fig S2:** Total collagen in circ-*crh-1* mutants at different temperatures Total collagen-to-protein ratio in *crh-1(syb385)* mutants compared to wild-type for 1-day adult worms at 15°C, 20°C, and 25°C. There is a non-significant difference in mean total collagen level between *crh-1*(*syb385*) mutants and wild-type animals (15°C, *p*=0.423; 20°C, *p*=0.632; 25°C, *p*=0.250). n=3 independent total collagen determination assays.

**Table S1:** Strains used in this study.

**Table S2:** Oligonucleotide primers used in this study.

**Table S3:** Summary and statistics of lifespan experiments.

## MATERIAL AND METHODS

### Strains, general animal cultivation and genetic controls

Worms were cultivated on the surface of NGM agar seeded with the *Escherichia coli* strain OP50 as the primary food source and grown in 20°C incubators using standard protocols unless indicated otherwise. All experiments were performed on hermaphrodites. The wild-type strain N2, variety Bristol (Brenner, 1974) and other strains used in this study are listed in **Table S1**. Strains were constructed using standard genetic methods (Fay, 2006) and genotypes were confirmed either by phenotype (for example, the transgenic strain was marked by fluorescence) or by PCR (for example, by identifying small deletions in mutant strains).

### Generation of plasmids, transgenic animals, and *col-49* mutants

To generate transgenic worms expressing the circ-*crh-1* in muscle cells, exon 4 of *crh-1* and intronic sequences flanking exon 4 (**Figure 4A**) were cloned into the pMC10 plasmid (a kind gift from the Sengupta Lab). Next, promoter sequences of *myo-3* (∼2.5 kb) were cloned at the 5’-end of the circ-*crh-1* sequence using the multi cloning site (MCS) of pMC10. The generated *myo-3p::circ-crh-1* construct along with the *unc-122p*::RFP co-injection marker (AddGene) were injected into VDL1300 *crh-1(syb385); dvIs100*[*unc-54p::A::unc-54 3’-UTR, mtl-2p::GFP*] animals to create VD12 (**Table S1**).

Transgenic worms carrying extrachromosomal arrays overexpressing *pie-1p::circ-crh-1* (VDL975), *rab-3p::circ-crh-1* (VDL1104) were crossed with the VDL1300 strain to create VDL1306 and VDL1307 strains (**Table S1**). A *col-49* mutant allele (*syb8747*) harboring a 1180bp deletion was generated using a Co-CRISPR method and confirmed by PCR and Sanger sequencing (SunyBiotech). sgRNAs used to generate the *col-49(syb8747)* mutant were Sg1: 5’-cctcatcatcatgtggaaattcg and Sg2: 5’-cccacctagaactgcttgattcg.

### Lifespan analysis

All strains were maintained at 20°C for at least two generations before the lifespan assay. Adult worms age-synchronized by hypochlorite treatment and collected eggs were hatched overnight at 20°C. L1 larvae were then plated onto NGM plates seeded with *E. coli* OP50 bacteria. At the L4 larval stage, 90-150 worms per genotype were transferred to new 6 cm NGM plates seeded with 10x concentrated *E. coli* OP50 bacteria containing 0.5 µM 5-fluorodeoxyuridine (FUdR) to inhibit the development of self-progeny, and then shifted at the young adult stage to 25°C. Each strain was assayed in parallel and each plate contained 10-15 worms. Worms were blindly scored every day and were considered dead when they did not respond to touch of the platinum wire pick and were subsequently removed from the plate. Worms that experienced ventral rupture, bagging, or walling were censored from the lifespan analysis.

### Paralysis assays

The paralysis assay was performed using GMC101 transgenic animals expressing *unc-54p::A*β*_1-42_* as described previously (McColl et al., 2012). Briefly, worms were age-synchronized by hypochlorite bleaching and cultivated at 20°C. After they reached the L4 larval stage, worms were transferred to assay plates freshly seeded with *E. coli* OP50 bacteria, containing 0.5 µM of FUdR to inhibit the development of self-progeny, and then shifted at the young adult stage to 25°C to induce paralysis. About 15 worms were placed on each 6 cm NGM plate, and animals were blindly scored every 24 hours as “paralyzed” if they failed to perform a full body wave propagation following a repeated touch-provoked response.

### Total RNA collection and extraction

Worms were age-synchronized worms by hypochlorite treatment and collected eggs were hatched overnight at 20°C in 1x M9 buffer. L1 larvae were then plated onto NGM plates seeded 10x concentrated *E. coli* OP50 bacteria and allowed to develop to the L4 larval stage at 20°C. L4 larvae were then collected, washed and re-plated onto *E. coli* OP50 seeded NGM plates containing 0.5 µM FUdR. Worms were either upshifted to 25°C or kept at 20°C. Adult worms were collected at different aging time-points and washed with 1x M9 buffer through 35 µM nylon mesh to remove bacteria. Worm pellets of 100-300 µl were then transferred into green RINO tubes (Next Advance) and TRizol LS reagent (ThermoFisher Scientific, Cat #10296028) was added in a 1:3 ratio. Worms were immediately lysed by bead beating them for 5 min using a Bullet Blender Pro Storm (Next Advance). Total RNA was extracted using the Purelink RNA mini-kit, followed by a DNAse I treatment following the manufacturer’s protocol (Ambion, Cat #12183020). RNA was quantified by a Nanodrop. Bioanalyzer or tapestation (Agilent) were used for qualification as needed. and samples were stored at −80°C.

### Analysis by RT-qPCR

To quantify and confirm individual circular or linear transcripts, 0.5 μg total RNA was reverse transcribed using Superscript III to prepare cDNA using random hexamers (Invitrogen, Cat #18080051). Next, cDNA samples were diluted and used with PowerUp SYBR Green Master Mix (Applied Biosystems, Cat #A25471) for RT-qPCR analysis analyzed on a CFX96 Real-Time System (Bio-Rad). For RT-qPCRs of circRNAs, we used outward-facing primers. For host gene linear RNA counterparts, one primer was located in the circularizing exon and the other was located in the upstream or downstream non-circularizing exon. For linear mRNAs such as collagen-encoding genes and *A*β mRNA, we used forward-facing primers. Fold-change values were calculated using wild-type (N2) ΔCt as control values for the 2^-ΔΔCt^ method. Data is normalized to housekeeping genes (*cdc-42* or *act-1*) mRNA. Primer sequences are listed in **Table S2**.

### Imaging and quantification of Aβ aggregates in living animals

Four-day old adults were randomly collected during paralysis assays and stained with 1 mM X-34 (Sigma, Cat # SML1954) 10 mM Tris-HCl pH 8.0 for 2 hours as previously described (Link et al., 2001a). Stained worms were then washed twice with 1x M9 and transferred to *E. coli* seeded NGM plates containing 0.5 µM FUdR for 24 hours to de-stain worms. De-stained 5-day old adults were then placed onto a 2% agarose pad with 10 µM levamisole to anesthetize worms. Confocal microscopy was used to acquire and capture images of the worm head using a 40x oil objective (405 nm excitation, 470-520 nm emission range). ImageJ software (NIH) was used to quantify Aβ aggregates.

### Total collagen level determination

As previously described (Teuscher et al., 2019), worms were age-synchronized by hypochlorite treatment and collected eggs were hatched overnight at 15°C, 20°C, or 25°C in 1x M9 buffer. L1 larvae were then plated onto NGM plates seeded *E. coli* OP50 bacteria and allowed to develop 24 hours post-L4 larval stage (1-day adult) at 15°C, 20°C, or 25°C. 1-day adult worms were then collected with 1x M9 buffer and washed with dH_2_O through a 35 µM nylon mesh to remove bacteria. Worm pellets of 300 µl were then transferred into green RINO tubes (Next Advance) and lysed by bead beating them for 10 min using a Bullet Blender Pro Storm (Next Advance). Total collagen level was determined using the QuickZyme Total Collagen Kit (QuickZyme Biosciences), following the manufacturer’s protocol. Briefly, lysate samples were mixed with 12M HCL solution and incubated for 20 hours at 95°C. Then assay buffer was added, and the 96-well plate was incubated at room temperature for 20 min, followed by the addition of the detecting reagent and incubation at 60°C for 60 min. Total collagen level was measured and quantified as a fraction of total protein abundance using a Synergy HT BioTek microplate reader. Total protein levels were quantified using the Pierce^TM^ BCA Protein Assay Kit (ThermoFisher Scientific) following the manufacturer’s protocol.

### Statistical analysis

Statistical comparisons and graphical representations were performed with the Online Application for Survival Analysis, OASIS 2 (Han et al., 2016). For lifespan survival curves, we used the Mantel-Cox log-rank test. Other data were analyzed using Graphpad Prism 9 software and statistical comparisons made include the Mann-Whitney *t*-test or the one-way ANOVA followed by a posthoc multiple-comparisons test. *p*-values are reported in the figure legends.

