## Supplementary figures and images for "Loss of age-accumulated *crh-1* circRNAs ameliorate amyloid β-induced toxicity in a *C. elegans* model for Alzheimer’s disease"

### Supplemental Figure 1

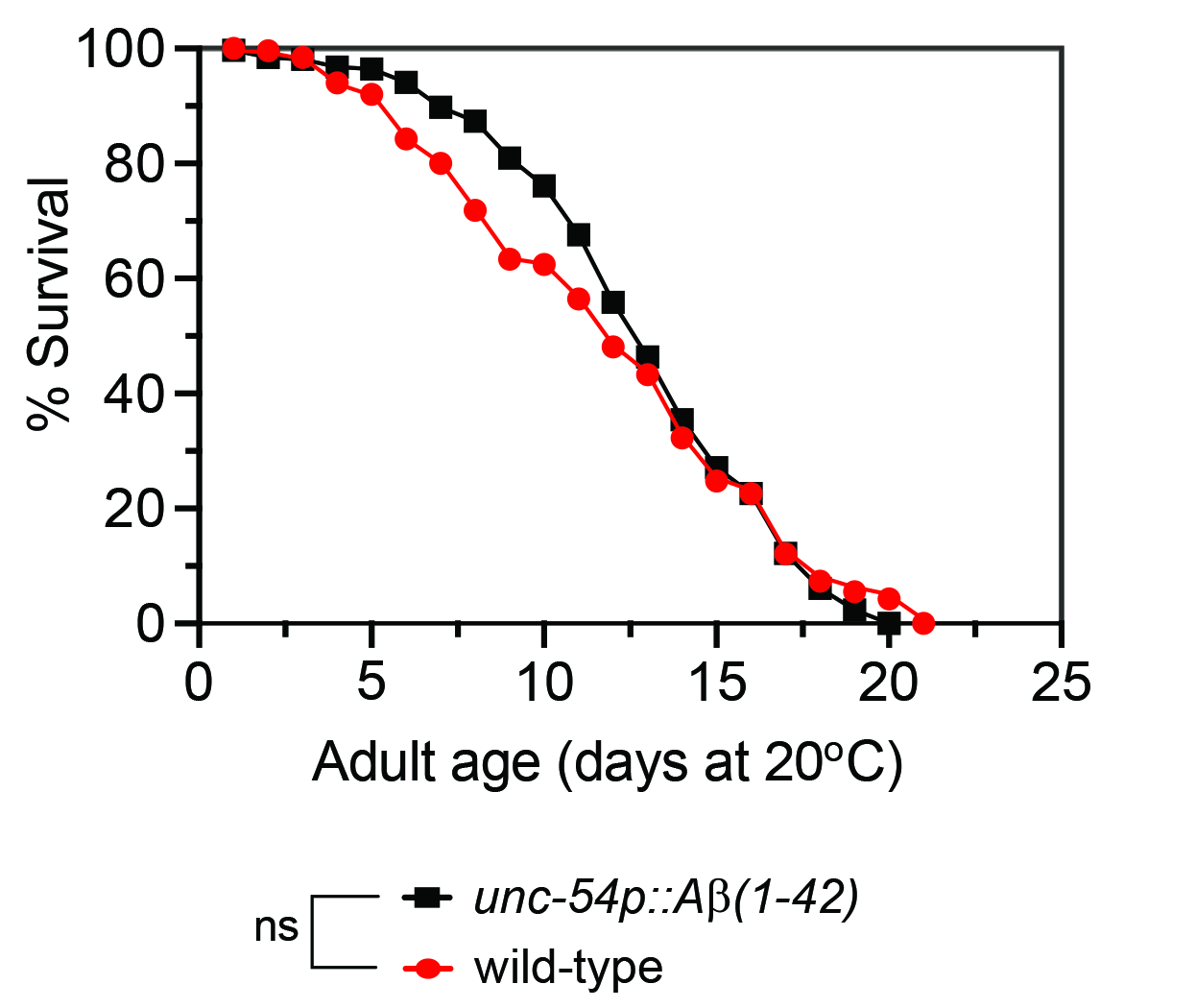

### Supplemental Figure 2

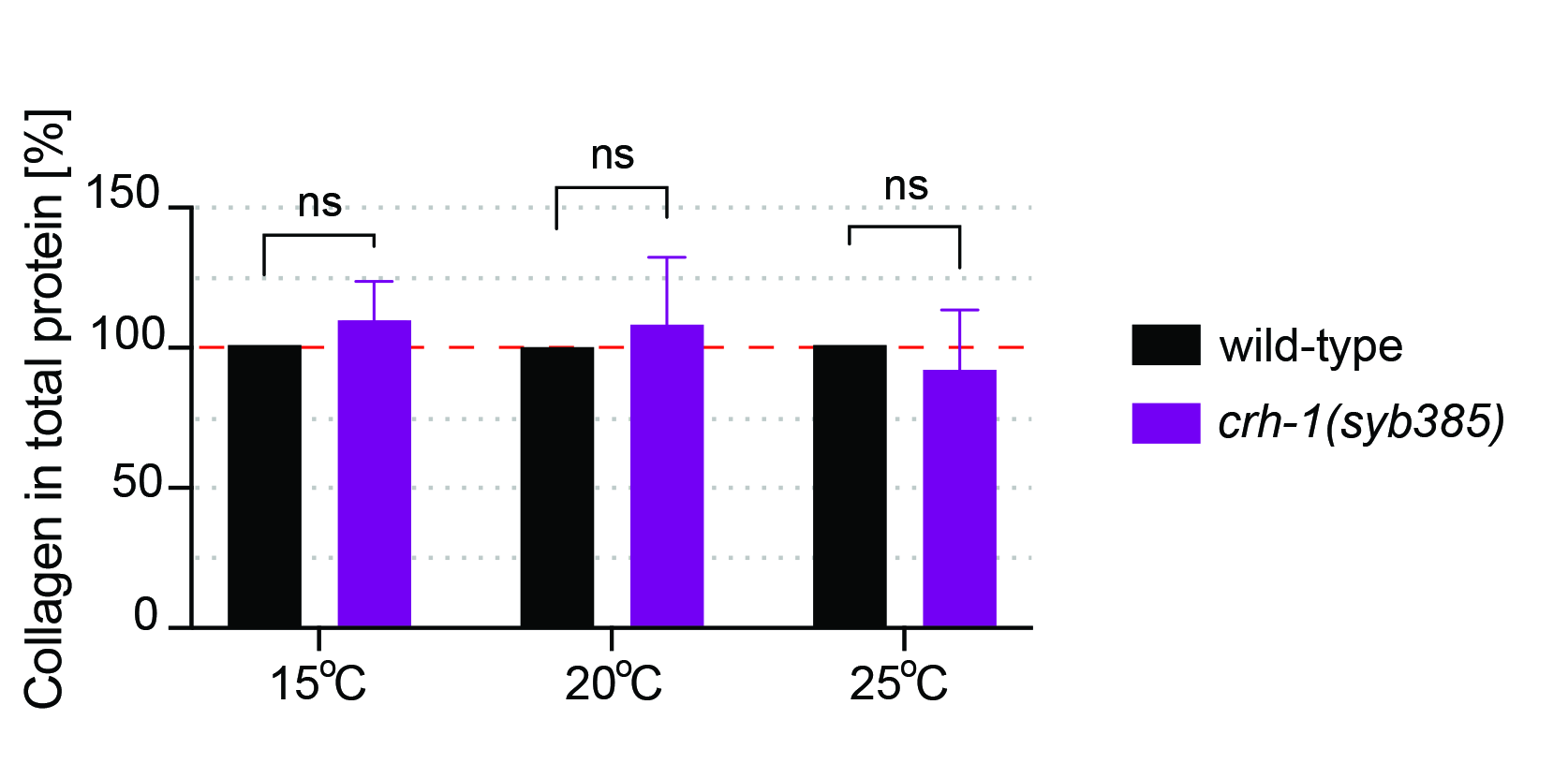
